# Norstictic acid is a selective allosteric transcriptional regulator

**DOI:** 10.1101/2021.03.26.437253

**Authors:** Julie M. Garlick, Steven M. Sturlis, Paul A. Bruno, Joel A. Yates, Amanda L. Peiffer, Yejun Liu, Laura Goo, LiWei Bao, Samantha N. De Salle, Giselle Tamayo-Castillo, Charles L. Brooks, Sofia D. Merajver, Anna K. Mapp

## Abstract

Inhibitors of transcriptional protein-protein interactions (PPIs) have high value both as tools and for therapeutic applications. The PPI network mediated by the transcriptional coactivator Med25, for example, regulates stress-response and motility pathways and dysregulation of the PPI networks contributes to oncogenesis and metastasis. The canonical transcription factor binding sites within Med25 are large (~900 Å^2^) and have little topology, and thus do not present an array of attractive small-molecule binding sites for inhibitor discovery. Here we demonstrate that the depsidone natural product norstictic acid functions through an alternative binding site to block Med25-transcriptional activator PPIs *in vitro* and in cell culture. Norstictic acid targets a binding site comprised of a highly dynamic loop flanking one canonical binding surface and in doing so, it both orthosterically and allosterically alters Med25-driven transcription in a patient-derived model of triple negative breast cancer. These results highlight the potential of Med25 as a therapeutic target as well as the inhibitor discovery opportunities presented by structurally dynamic loops within otherwise challenging proteins.

## Introduction

Transcriptional coactivators play an integral role in the regulation of gene expression, serving as hub proteins for transcriptional machinery assembly through interactions with transcriptional activators.^1–9^ Alterations in the network of coactivator-activator protein-protein interactions (PPIs) contribute to the onset and perpetuation of numerous diseases, leading to significant interest in synthetic probes for mechanistic studies and therapeutic applications.^10–16^ This is especially true for coactivators whose function is highly context-dependent, required only for a subset of genes or only at particular times or locations in the life cycle of an organism.^17^ Thus, synthetic modulation of such coactivators would also be contextdependent, providing an advantageous layer of specificity. Recent structural and functional studies of the coactivator Med25 indicate that it falls into this category (Figure 1A); homozygous deletion of Med25, for example, is nonlethal, impacting approximately 900 genes.^18^ Several lines of evidence indicate that dysregulation of the PPI network of Med25 and the ETV/PEA3 transcriptional activators contributes to oncogenesis as well as metastatic phenotypes in certain breast and prostate cancers, heightening a need for Med25-selective inhibitors.^19–22^ Here we report the discovery of the natural product norstictic acid as the first such inhibitor.

**Figure 1.**
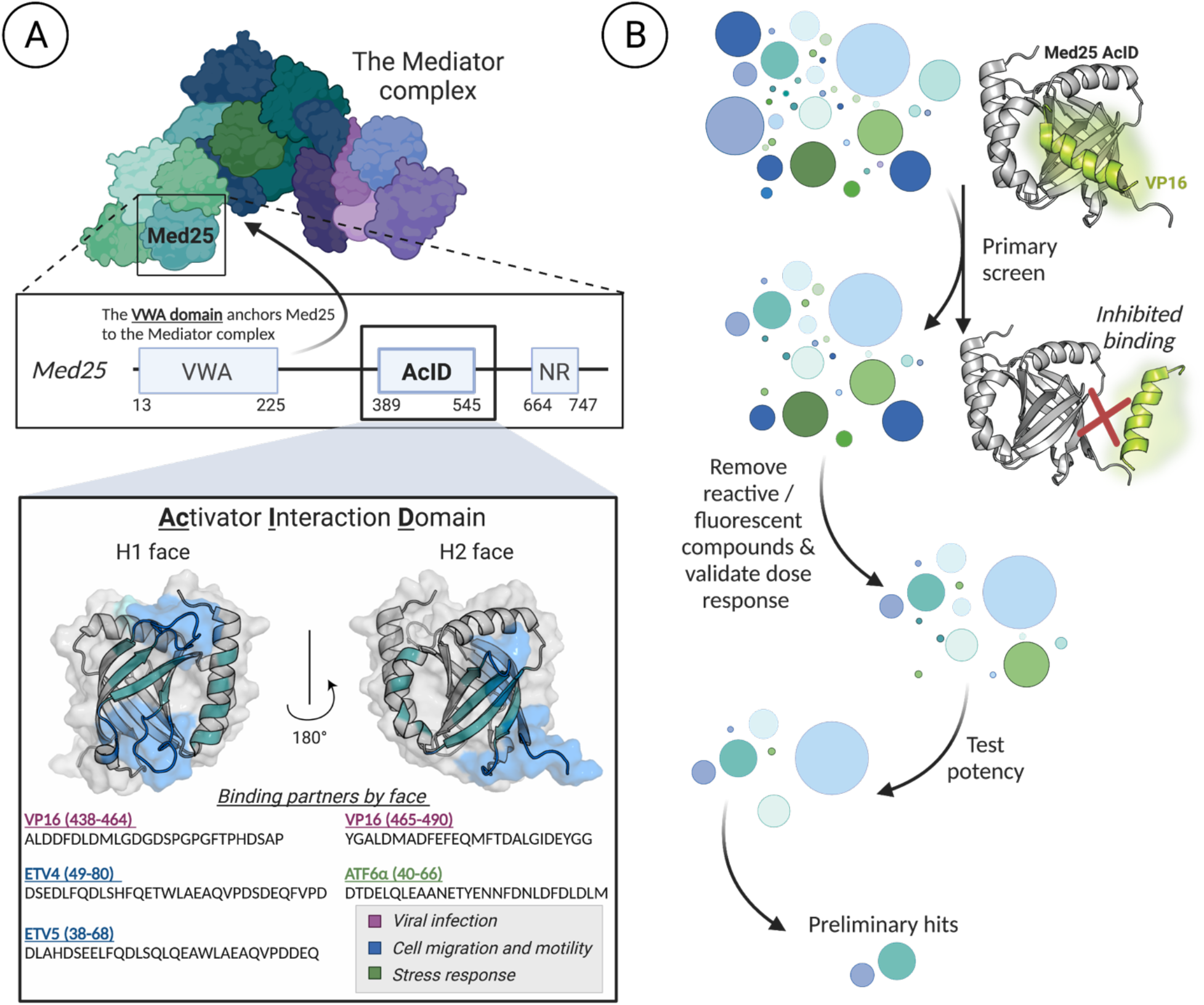
A) Hub protein Med25 is a subunit of the Mediator coactivator complex. The <u>Ac</u>tivator-<u>I</u>nteraction <u>D</u>omain (AcID) forms a PPI network with transcriptional activators using two binding surfaces (H1, H2) and in doing so regulates key cellular processes.^19,22-25^ PDB 2XNF used to generate figure. B) Schematic of HTS screen to identify inhibitors of the Med25 AcID PPI network. See SI for full screening details.

The domain of Med25 that interacts with activators is the <u>Ac</u>tivator-<u>I</u>nteraction <u>D</u>omain (AcID) (Figure 1A).^19,22-28^ This domain contains a 7-stranded beta barrel core flanked with dynamic loops and three alpha helices. Like other activator-binding domains (ABDs) within coactivators, AcID does not contain defined binding pockets, but rather relies on hydrophobic interfaces, termed H1 and H2, to interact with transcription factors.^23,25^ These qualities make AcID as well as other coactivators, challenging to target selectively, particularly in an orthosteric mode.^17,29^ Recently, we reported that the dynamic substructures within AcID contain allosteric hotspots that regulate cooperativity and selectivity in binding.^20,28^ Thus, we hypothesized that these substructures, largely loop regions, represent an opportunity for allosteric inhibition. Further, because such substructures are more likely to access conformations with topologically unique binding surfaces, one might anticipate that small-molecule modulators of such sites would exhibit enhanced selectivity compared to purely orthosteric ligands.^12,30-34^

## Results and Discussion

To identify inhibitors of Med25 AcID, we utilized a high-throughput fluorescence polarization (FP) assay interrogating a complex of AcID and fluorescein-tagged VP16(465-490) (Figure 1B, SI Figure S1). As previously reported, this VP16 sequence contains the minimal binding sequence for interaction with AcID (K_D_ = 0.60 ± 0.06 μM) and interacts with the H1 and H2 binding surfaces.^20,23^ Several commercially available libraries (MS Spectrum 2000, Focused Collections, and BioFocus NCC libraries) with a combined total of 4046 compounds were screened using this format (Z’ = 0.87; 1.6% hit rate; see SI for additional details). Compounds with activity >3 S.D. relative to the negative control (DMSO) were subjected to dose-response assessment with freshly purchased material, as well as secondary selectivity assays. From this, the lichen-derived natural products norstictic acid (NA) and psoromic acid (PA) emerged as the best inhibitors, with apparent IC_50_ values of 2.3 ± 0.1 μM and 3.9 ± 0.3 μM, respectively (Figure 2A,B).

**Figure 2.**
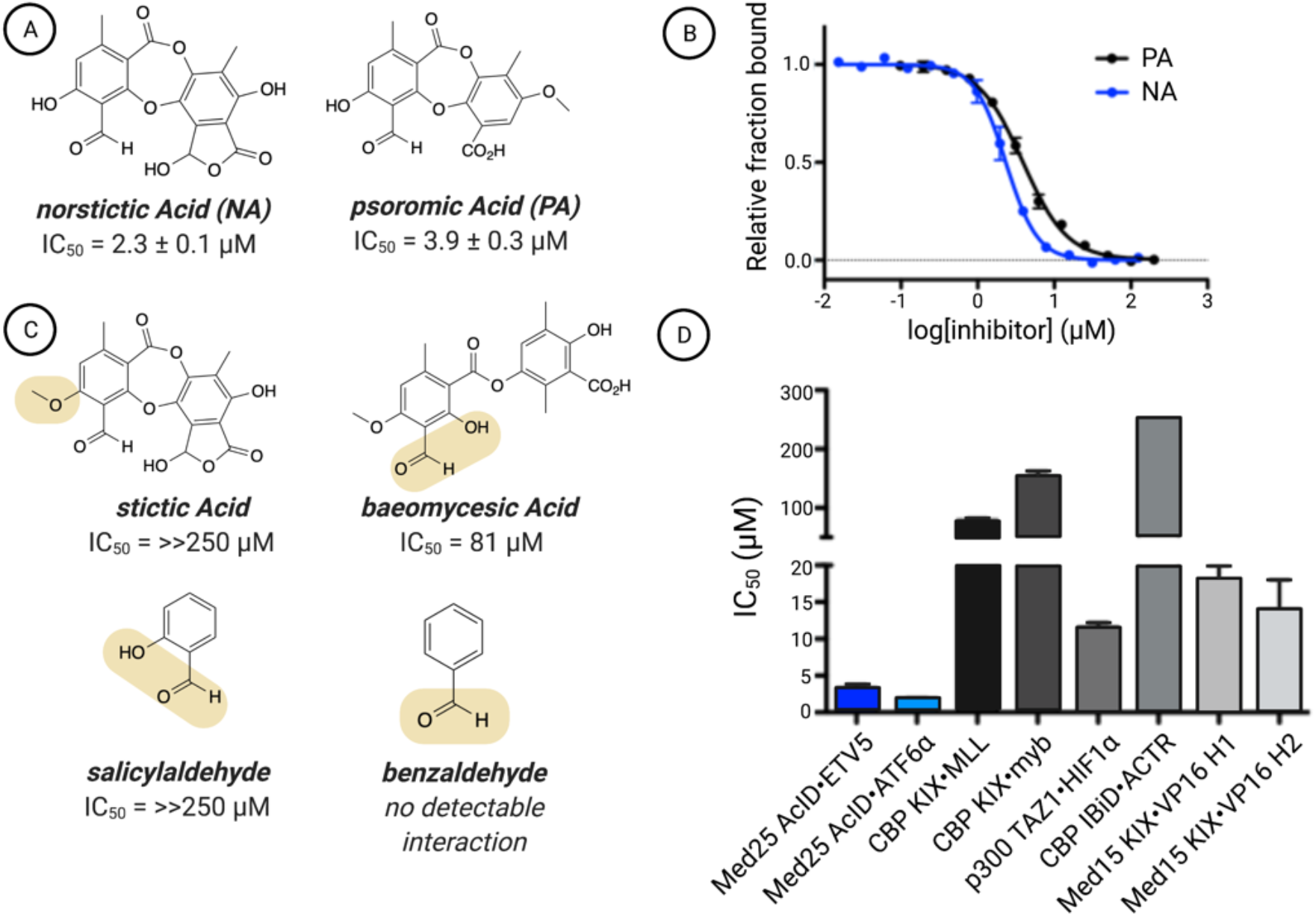
A) Chemical structures of the top two hits emerging from the screen of the Med25 AcID•fl-VP16(465-490) along with B) their apparent IC_50_ values. The apparent IC_50_ values were determined through titrations of either NA or PA against Med25 AcID•fl-VP16(465-490) performed in triplicate with the indicated error (SDOM). Full experimental details reported in the Supporting Information. C) Assessment of related structures shows that the orthophenoxyaldehyde moiety is important but not sufficient for inhibitory activity. IC_50_ values were determined via competition fluorescence polarization against Med25•VP16(465-490). D) Inhibition of related PPI networks by NA. Apparent IC_50_ values were measured via fluorescence polarization against a suite of coactivator domains (CBP KIX, p300 TAZ1, CBP IBiD, Med15 KIX) bound to fluorescein-tagged activators. The values are the average of three independent experiments with the indicated error (SDOM). No error bars are shown for the IC_50_ against IBiD•ACTR because the IC_50_ was greater than the highest concentration of NA tested, 250 μM, and thus we can only accurately report the IC50 as > 250 μM..Full details in the Supporting information.

Both NA and PA are natural products in the depsidone family containing an orthophenolic aldehyde moiety. The presence of a reactive aldehyde functionality suggested a potential covalent mechanism of action, for example via imine formation with lysine side chains. Consistent with this hypothesis, analysis of NA-treated Med25 AcID using mass spectrometry showed the presence of concentration-dependent covalent adduct(s) (SI Figure S2). Treatment with the reducing agent NaBH_4_ led to incorporation of H_2_ into the adduct, indicating initial formation of a Schiff base followed by reduction (SI Figure S3). Data from a time-course experiment revealed that at 5 minutes, significant inhibition is observed, with full activity after 30 minutes (SI Figure S4). An examination of related structures indicates that the orthophenolic aldehyde is necessary, but not sufficient for interaction with Med25 AcID or for inhibitory activity. Stictic acid, in which the phenol is masked as a methyl ether, inhibits Med25 interactions poorly (IC_50_ > 250 μM) (Figure 2C). Additionally, salicylaldehyde efficiently labels Med25 AcID, but does not impact binding of activators (SI Figure S5). These data suggest that noncovalent interactions play essential roles in the inhibitor function of NA. Consistent with this, NA exhibits remarkable selectivity for Med25 PPIs relative to other coactivators with similar binding surfaces (Figure 2D). Notably, NA inhibits Med25 PPIs at both binding surfaces, including those formed with transcriptional activators ETV5 (H1 binding surface) and ATF6α (H2 binding surface).

Several lines of evidence suggested that the engagement site of NA is a lysine-rich dynamic loop that borders the H2 binding surface (Figure 3A). There are 11 lysine residues within Med25 AcID, 6 of which are found on dynamic loop regions flanking the two known activator binding surfaces. Replacement of these lysines with arginine either alone or in combination had minimal effects on binding of the cognate transcriptional activator binding partners (Figure S6, S7). Similarly, mutations within the H1 binding surface had minimal impact on both NA binding, determined by mass spectrometric analysis, and inhibition in an *in vitro* binding assay (Figure 3A, B; SI Figure S8). In contrast, mutation of K519 had a profound effect on NA binding and inhibition. This residue is part of a lysine-rich dynamic loop that flanks the H2 face and the mutational data indicates that NA can also interact with K520 and K518 within this loop.

**Figure 3.**
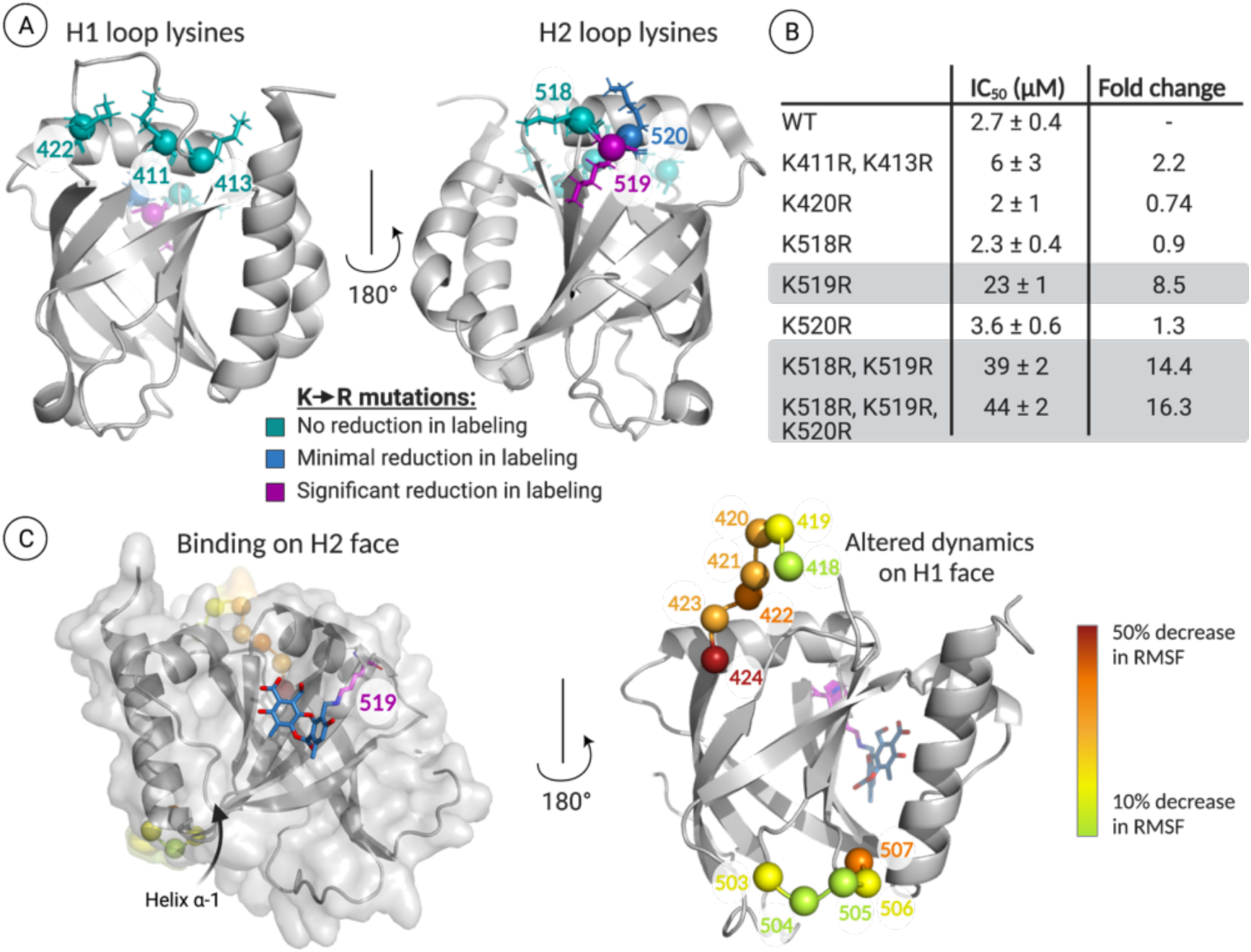
A) LC-MS analysis of norstictic acid covalent adduct formation with Med25 lysine-to-arginine mutants indicates that K519R leads to the most significant reduction of labelling. No reduction of labelling corresponds to a decrease in abundance of the NA covalent adduct of less than 10%. Minimal reduction in labelling, observed for K520R, corresponds to a 22% reduction in the mass abundance of the NA covalent adduct. Significant reduction in labelling, observed for K519R, corresponds to a 53% reduction in the mass abundance of the +1 covalent adduct. See Supplemental Information for additional quantitative analysis. PDB 2XNF used to generate figure. B) Inhibition of Med25 AcID•ETV5 interaction by norstictic acid measured using fluorescence polarization. Mutants containing K519R, highlighted in grey, demonstrate the most significant increase in apparent IC_50_. Values represent the average of three independent experiments with the indicated error (SDOM) C) (Left) Centroid structure of the most populated cluster from molecular dynamics simulations, where norstictic acid binds to the H2 face of Med25 and covalently links to K519. (Right) The residues that showed the greatest reduction in fluctuations (RMSF) upon activator binding all occur on dynamic substructures on the H1 face.

To develop a structural model of NA binding and function, molecular dynamics simulations of the covalent NA-Med25 AcID complex in which NA is covalently linked to K519 were carried out, and the results compared to the case of unbound Med25 AcID. As illustrated in Figure 3C, minimal restructuring in the lysine loop adjacent to the H2 binding interface is observed. However, helix a1 shows significant conformational changes, resulting in partial unfolding. More surprising, the only detectable dynamical changes in NA binding occur on the H1 face, with residues in the two loops on that face showing up to 50% reduction in root-mean-square fluctuations (RMSF). Taken together, the data indicate that NA serves as both an orthosteric inhibitor of H2-binding transcription factors (e.g. ATF6α) and an allosteric inhibitor of H1 binding transcriptional activators (e.g. ETV5).

Next, we tested the engagement of full-length Med25 by NA and the resulting impact of PPI formation and function. Thermal shift assays using freshly prepared HeLa nuclear extracts demonstrate that NA stabilizes endogenous Med25 protein, indicating engagement with the AcID motif in the context of fulllength protein (Figure 4A).^35,36^ The ability of NA to block PPIs formed between endogenous Med25 and cognate activators was then assessed. ETV5 is a member of the ETV/PEA3 subfamily of transcriptional activators, comprised of ETV5, ETV1, and ETV4. This transcriptional activator trio has nearly identical domains that utilize a PPI with the H1 surface of Med25 for function, and the PPIs are dysregulated in cancer through overexpression of one or both of the binding partners.^19,21,37^ As shown in Figure 4B, NA treatment of HeLa cells blocks formation of the Med25•ETV5 complex, consistent with the *in vitro* binding data of Figure 2D. The Med25•ETV/PEA3 PPIs regulate proliferation, invasion and migration pathways and in at least a subset of cancers, are part of a Her2-driven RAS-RAF-MEK-MAPK circuit..^38,39^ Thus, if NA blocks Med25•ETV/PEA3 PPIs, positive synergy with a Her2 inhibitor would be anticipated in such systems. To test this, the combination of NA and lapatinib was tested for synergy by the isobologram method (Figure 4C) in MDA-MB-231 breast cancer cells, an established model.^19^ As can be seen, strong synergy was observed, consistent with NA engagement of Med25, blocking PPIs formed with ETV/PEA3 activators. This model was further tested in the patient-derived early passage triple negative breast cancer cell line VARI068 with robust EGFR expression.^40,41^ VARI068 cells exhibit approximately 2-fold upregulation of Med25 relative to normal-like non-tumorigenic MCF10A cells. Treatment of VARI068 cells with NA blocks the Med25•ETV5 complex (Figure 4D). To further explore the effects of Med25 regulation on cancer, CRISPR-Cas9-mediated knockout of Med25 in VARI068 cells led to downregulation of Med25•ETV/PEA3-regulated MMP2 (Figure 4E) and NA treatment leads to substantial down-regulation of MMP2 relative to vehicle. Taken together, these data are consistent with a mechanism in which NA engages Med25 in cells and alters its PPI network with downstream effects on tumor phenotype.

**Figure 4.**
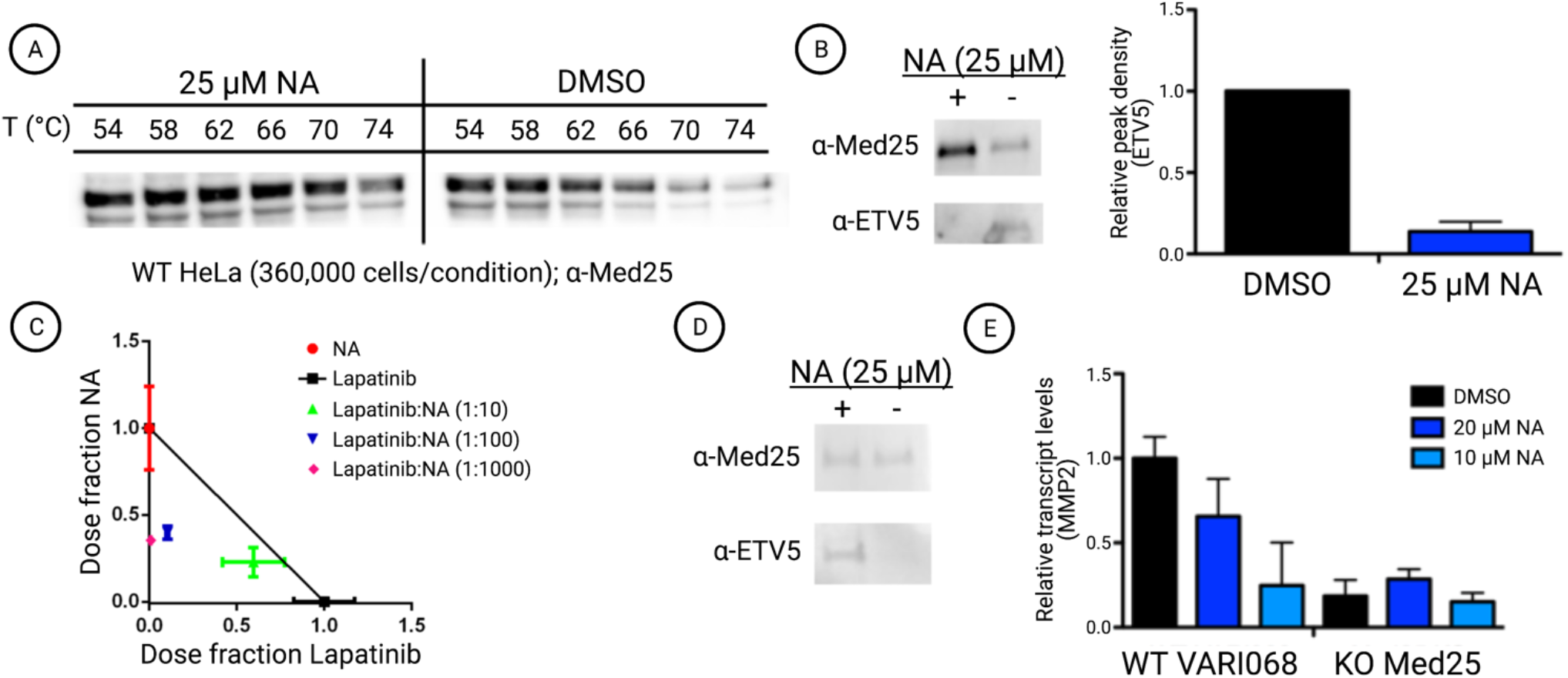
A) Norstictic acid significantly stabilizes full length Med25 in HeLa cell extracts. Cellular thermal shift assays were conducted by dosing HeLa cell nuclear extracts with 25 μM NA or equivalent DMSO and subjecting samples to a range of temperatures. Western blot using a Med25 antibody shows increased band density in NA dosed compared to control samples, indicating thermal stabilization and target engagement. Quantification and CETSA at additional concentrations available in the Supplemental Information, Figure S9. B) Treatment of HeLa cells with 25 μM NA attenuates the formation of the Med25-ETV5 complex. Left: A representative Western Blot shows a reduction in co-immuniprecipition of ETV5 with Med25. Right: Quantitative assessment of ETV5 Co-immunoprecipitation with Med25. Western blot band density was measured using ImageJ and normalized by comparison to overall Med25 levels. Results are the average of biological triplicate. See Supplemental information Figure S11 for blot images used. C) NA shows positive synergy with an on-pathway kinase inhibitor, lapatinib. IC_50_ values of fixed dose ratios of NA and lapatinib were measured in MDA-MB-231 cells after 2 days of dosing and plotted on an isobologram. D) Western blot shows that treatment of VARI068 cells with NA attenuates formation of the Med25-ETV5 complex. E) Analysis of MMP2 transcript levels by qPCR indicates that NA treatment decreases MMP2 levels to that of a KO variant of the cell line. MMP2 transcript levels are normalized to the reference gene RPL19. Results shown are the average of technical triplicate experiments, conducted in biological duplicate.

## Conclusions

The demonstration of NA as a selective orthosteric/allosteric inhibitor of Med25 function validates the importance of dynamic loops in coactivators in molecular recognition and their utility as targets. We also provide evidence that modulation of these protein-protein interactions would provide a useful tool to study cancer at the bench. Our work suggests that Med25 is a potentially viable therapeutic target, thus justifying the search for a drug-like compound for testing the clinical utility of the strategy. We anticipate that NA will be a useful tool for dissecting the Med25-ETV/PEA3 axis in cancers in which Med25 dysregulation is a hallmark. Further, the strategy of targeting dynamic substructures within coactivators should be generalizable beyond Med25.

## Acknowledgements

The authors acknowledge financial support from the National Institutes of Health (CA242018 to A.K.M.; NIH T32GM008597 to support S.D.S.; GM103695, GM130587 for C.L.B. III), the Breast Cancer Research Foundation (S.D.M, J.A.Y, L.W.B) and NIH-P30CA046592 (Rogel Cancer Center Core grant).

