## Supporting Information for "Norstictic acid is a selective allosteric transcriptional regulator"

**Table of Contents**

Figure S-1. Optimization of VP-16(465-490) binding assay for use in high throughput screening
Figure S-2. Mass spectrometry data showing concentration dependent norstictic acid adduct formation
Figure S-3. Treating with NaBH4 after incubation with NA confirms modification through initial imine formation
Figure S-4. Time course inhibition of VP16(465-490)•AcID by NA
Figure S-5. Mass spectrometry and competition FP with salicylaldehyde
Figure S-6. Direct binding curves of VP16(465-490) to various Med25 AcID lysine mutants
Figure S-7. Direct binding curves of ETV5(38-68) to various Med25 AcID lysine mutants
Figure S-8. Effect of lysine mutation on the covalent labeling of Med25 AcID by NA
Figure S-9. Root mean square fluctuations by Med25 AcID residue for the apo protein and the protein with NA bound
Figure S-10. NA thermally stabilizes Med25 in cell lysates at multiple concentrations
Figure S-11. Western blots showing NA inhibition of co-immunoprecipitation of ETV5 with Med25 in HeLA cells
Figure S-12. Dose-effect curves looking at MDA-MB-231 cell viability dosing with NA and lapatinb in various combinations
Figure S-13. Generation of Med25 CRISPR knock out cell line
Protein expression and purification
Synthesis of transcriptional activation domain peptides
Direct binding and competition fluorescence polarization
High-throughput screening
Mass spectrometry analysis of covalent adducts
Molecular dynamics simulations
Cell Culture
Thermal shift assays
Co-Immunoprecipitation
Quantitative polymerase chain reaction
Syngergy experiments with lapatinib
Generation of Med25 CRISPR KO cell lines
Analytical HPLC traces of synthetic peptides
References

### Supplementary Figures

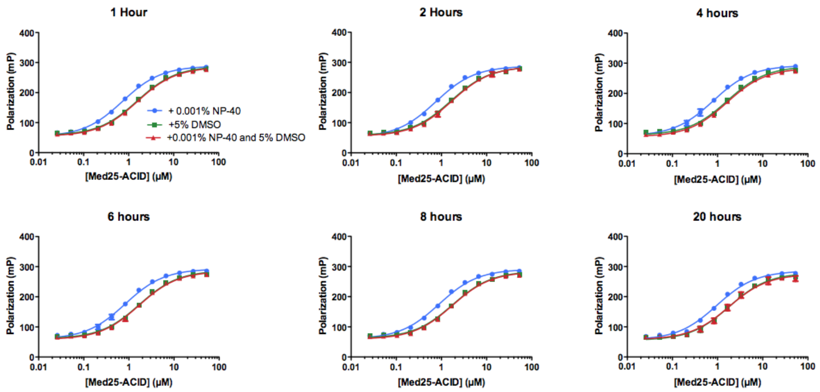


| | $K_d$ of Flo-VP16(465-490)-ACID interaction | | | | | |
|---|---|---|---|---|---|---|
| | 1 hour | 2 hour | 4 hour | 6 hour | 8 hour | 20 hour |
| + NP-40 | 0.72 µM | 0.75 µM | 0.80 µM | 0.81 µM | 0.88 µM | 0.95 µM |
| +DMSO | 1.5 µM | 1.6 µM | 1.6 µM | 1.7 µM | 1.7 µM | 1.8 µM |
| + NP-40 and DMSO | 1.5 µM | 1.7 µM | 1.6 µM | 1.7 µM | 1.7 µM | 1.8 µM |

**Figure S1**. Optimization of VP-16(465-490) binding assay for use in high throughput screening. Assay stability time course, DMSO, and NP-40 effects on affinity are shown. The effects of 0.001% NP-40 and 5% DMSO are shown in isolation (blue and green curves, respectively) or in combination (red curve). Curves represent the means of three independent experiments with error bars representing the standard deviation of the anisotropy at the indicated concentration of AcID protein.

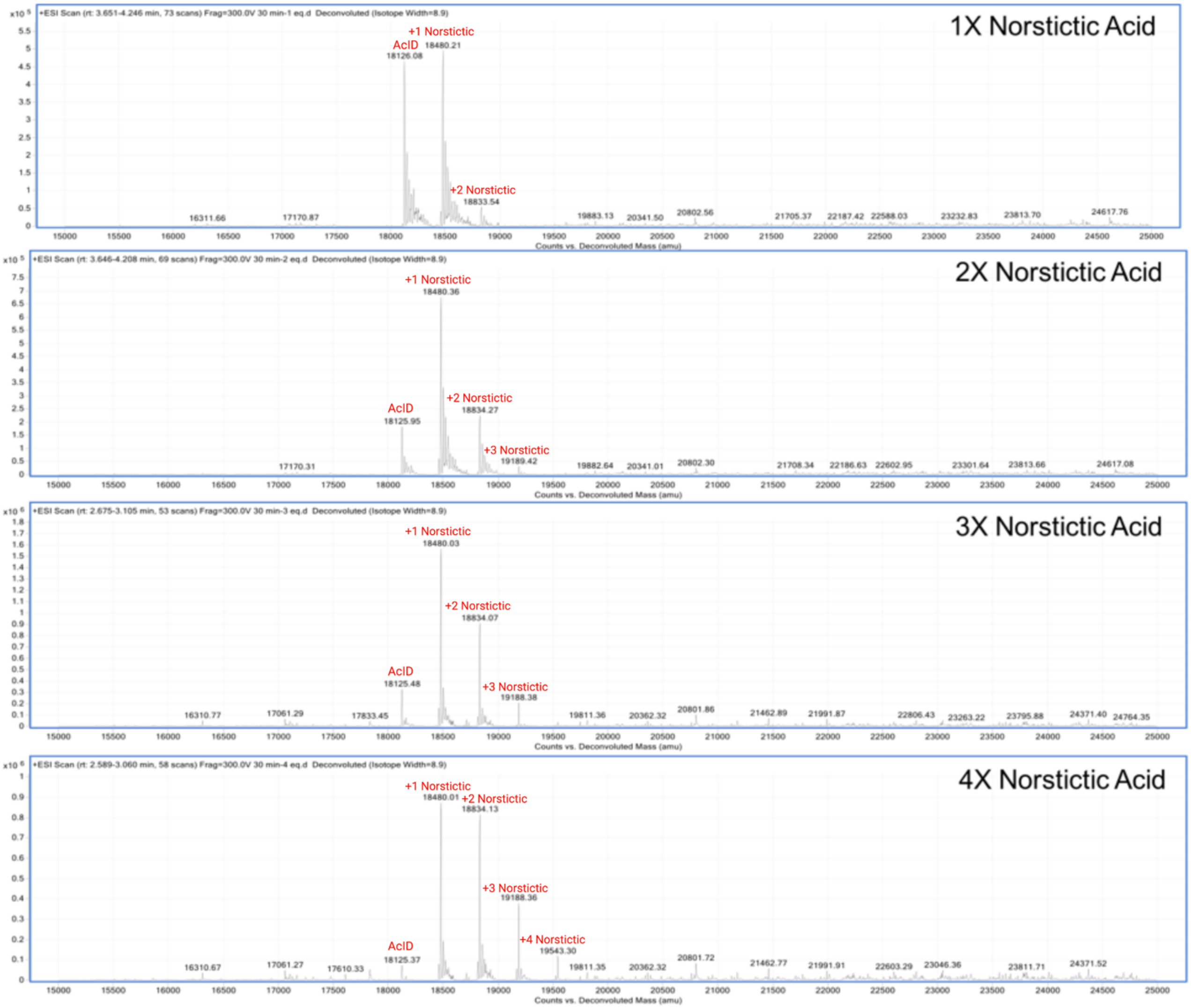


**Figure S2.** Concentration dependent norstictic acid adduct formation is observed with Med25 AcID (20 µM) using Mass Spectrometry. The NA modification corresponds to a mass increase of 354 Da. Data was obtained after incubation of Med25 with NA for 30 minutes.

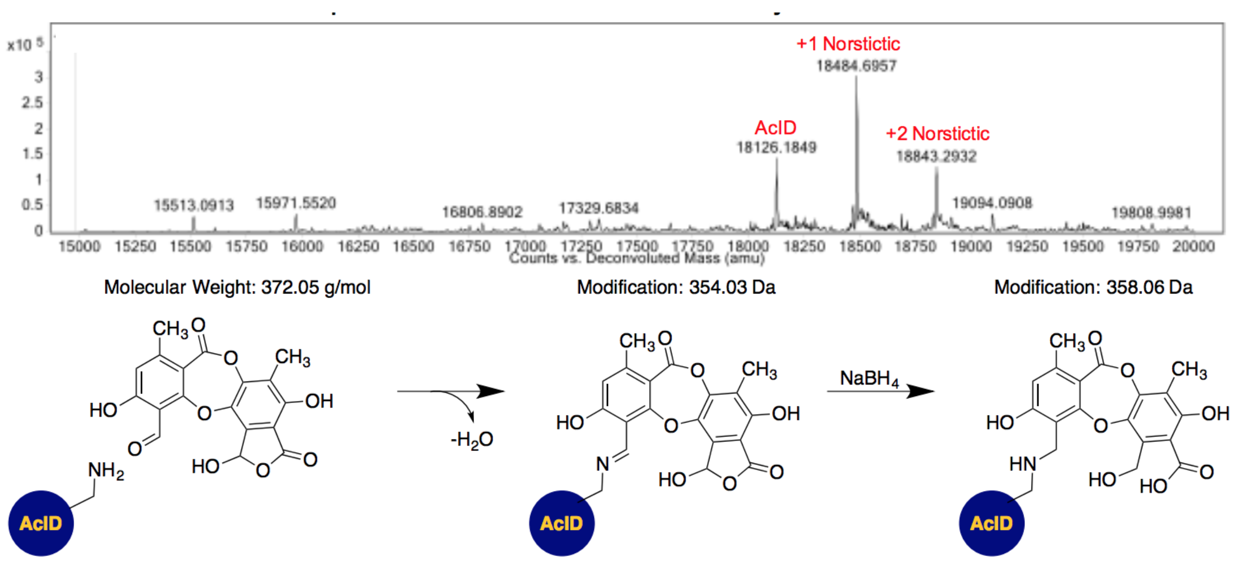


**Figure S3.** Addition of 4 equivalents $NaBH_4$ after incubation with NA for 30 minutes leads to formation of a +358 adduct (top), indicating covalent modification through initial imine formation (bottom).

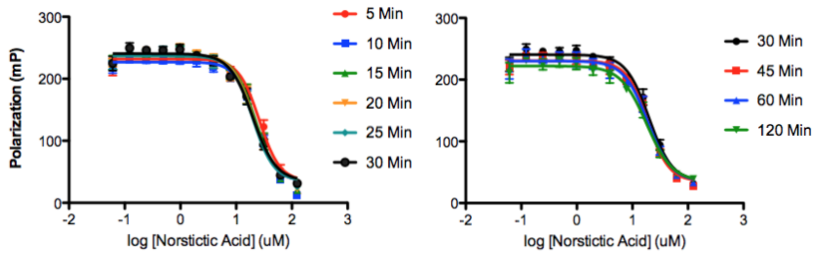


| | $IC_{50,app}$ (μM) |
|---|---|
| **5 min** | 26 ± 3 |
| **10 min** | 22 ± 2 |
| **15 min** | 22 ± 2 |
| **20 min** | 21 ± 2 |
| **25 min** | 20 ± 2 |
| **30 min** | 19 ± 2 |
| **45 min** | 19 ± 2 |
| **60 min** | 19 ± 2 |
| **120 min** | 18 ± 2 |

**Figure S4.** Time course inhibition of VP16(465-490)•AcID by NA. The apparent $IC_{50}$ of NA was determined for inhibition of the VP16(465-490)•AcID interaction at various time points to determine how quickly inhibition was achieved. Curves represent the mean values of three independent experiments, with vertical error bars representing the standard deviation of the mean polarization at the indicated concentration of norstictic acid.

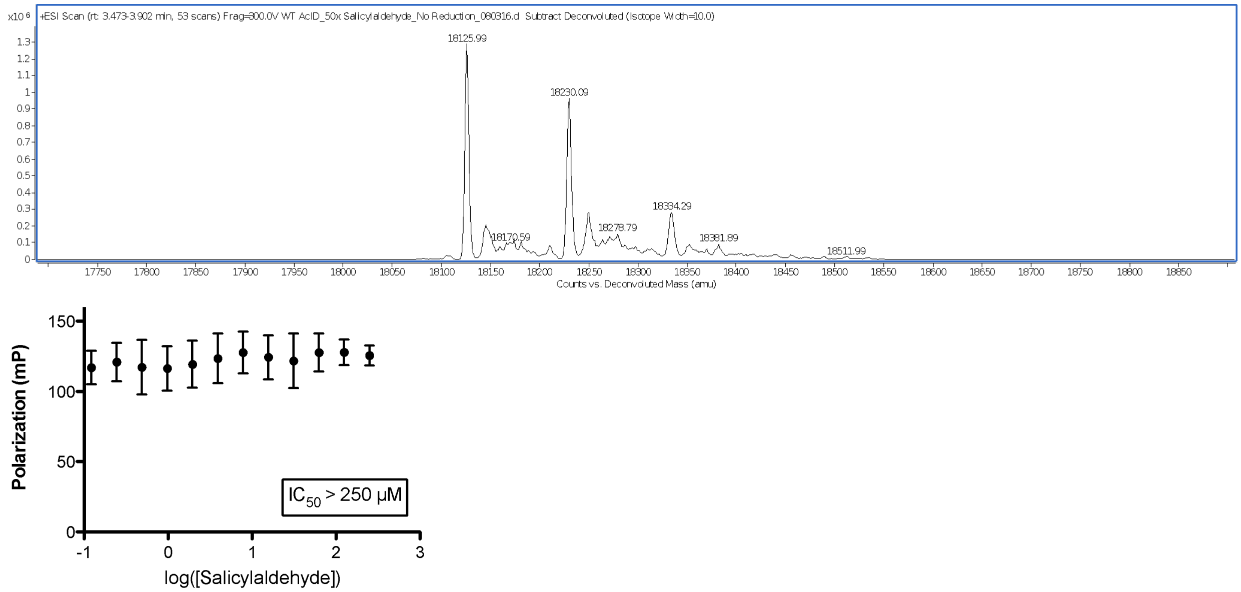


**Figure S5**. Salicylaldehyde covalently modifies, but does not inhibit, Med25 AcID. Mass spectrometry shows dosing with salicylaldehyde leads to formation of +104 Med25 adduct (top). Salicylaldehyde does not inhibit the Med25•VP16(465-490) interaction using fluorescence polarization (bottom). Curves represent the mean values of three independent experiments with vertical error bars representing the standard deviation of the fraction of tracer bound at the indicated AcID concentration.

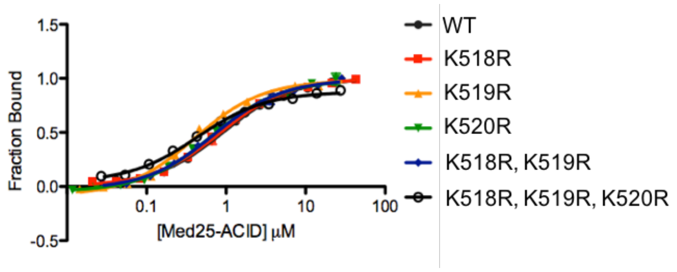


| | $K_D$ (μM) |
|---|---|
| **WT** | 0.90 ± 0.08 |
| **K518R** | 0.84 ± 0.08 |
| **K519R** | 0.39 ± 0.05 |
| **K520R** | 0.65 ± 0.08 |
| **K518R, K519R** | 0.69 ± 0.08 |
| **K518R, K519R, K520R** | 0.43 ± 0.05 |

| | $IC_{50}$ (μM) | Fold change |
|---|---|---|
| **WT** | 7.3 ± 0.9 | - |
| **K518R** | 8 ± 1 | 1.1 |
| **K519R** | 18 ± 5 | 2.5 |
| **K520R** | 13 ± 2 | 1.8 |
| **K518R, K519R** | 26 ± 5 | 3.5 |
| **K518R, K519R, K520R** | 35 ± 9 | 4.8 |

**Figure S6.** Direct binding curves of VP16(465-490) to various Med25 AcID lysine mutants (top). Apparent $IC_{50}$ values and fold changes associated with norstictic acid inhibition of VP16(465-490) interaction with mutant AcID constructs (bottom). Curves represent the mean values of three independent experiments with vertical error bars representing the standard deviation of the fraction of tracer bound at the indicated AcID concentration.

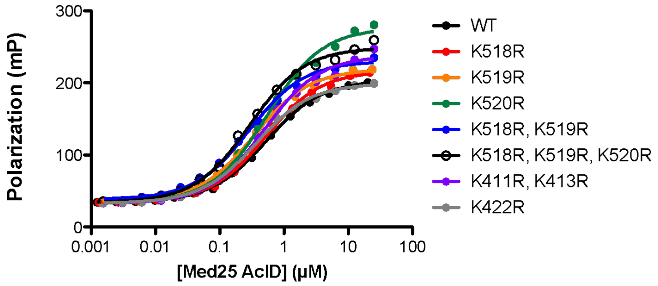


| | $K_D$ (μM) |
|---|---|
| **WT** | 0.52 ± 0.01 |
| **K518R** | 0.51 ± 0.01 |
| **K519R** | 0.34 ± 0.01 |
| **K520R** | 0.54 ± 0.02 |
| **K518R, K519R** | 0.28 ± 0.01 |
| **K518R, K519R, K520R** | 0.29 ± 0.01 |
| **K411R, K413R** | 0.49 ± 0.03 |
| **K422R** | 0.36 ± 0.01 |

**Figure S7**. Direct binding of ETV5(38-68) to Med25 AcID lysine mutants. Curves represent the mean values of three independent experiments with vertical error bars representing the standard deviation of the fraction of tracer bound at the indicated AcID concentration.

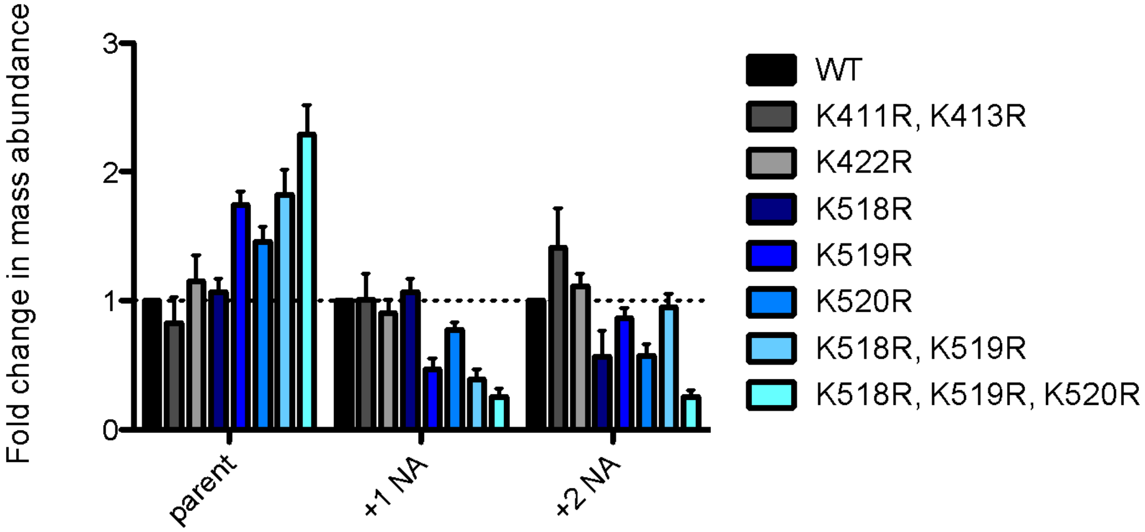


**Figure S8.** Effect of lysine mutation on the covalent labeling of Med25 AcID by NA. Protein was incubated with 1X NA for 30 minutes before conducting LC-MS analysis. Abundance of the parent mass and NA covalent adducts was determined and fold change compared to WT was calculated.

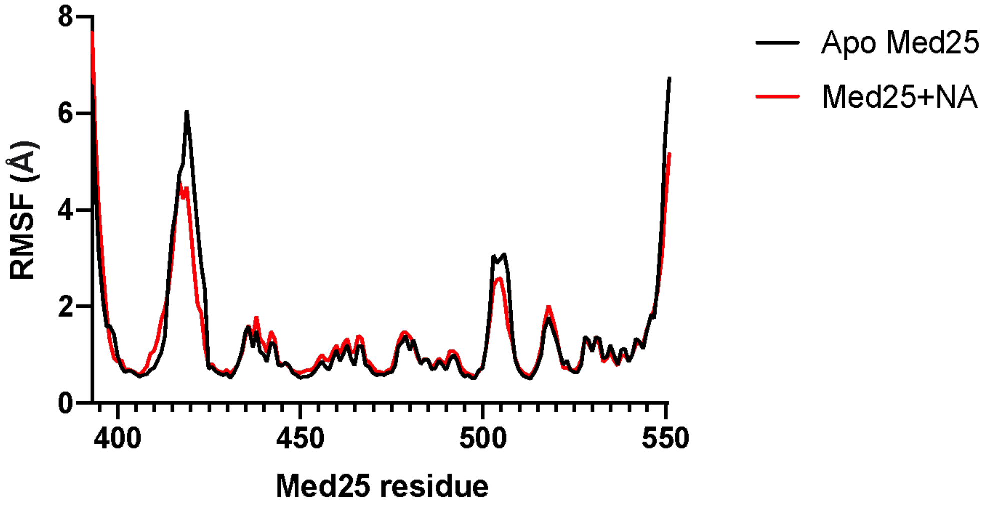


**Figure S9.** Root mean square fluctuations (RMSF) by Med25 AcID residue for the apo protein (black) and the protein with NA bound (red).

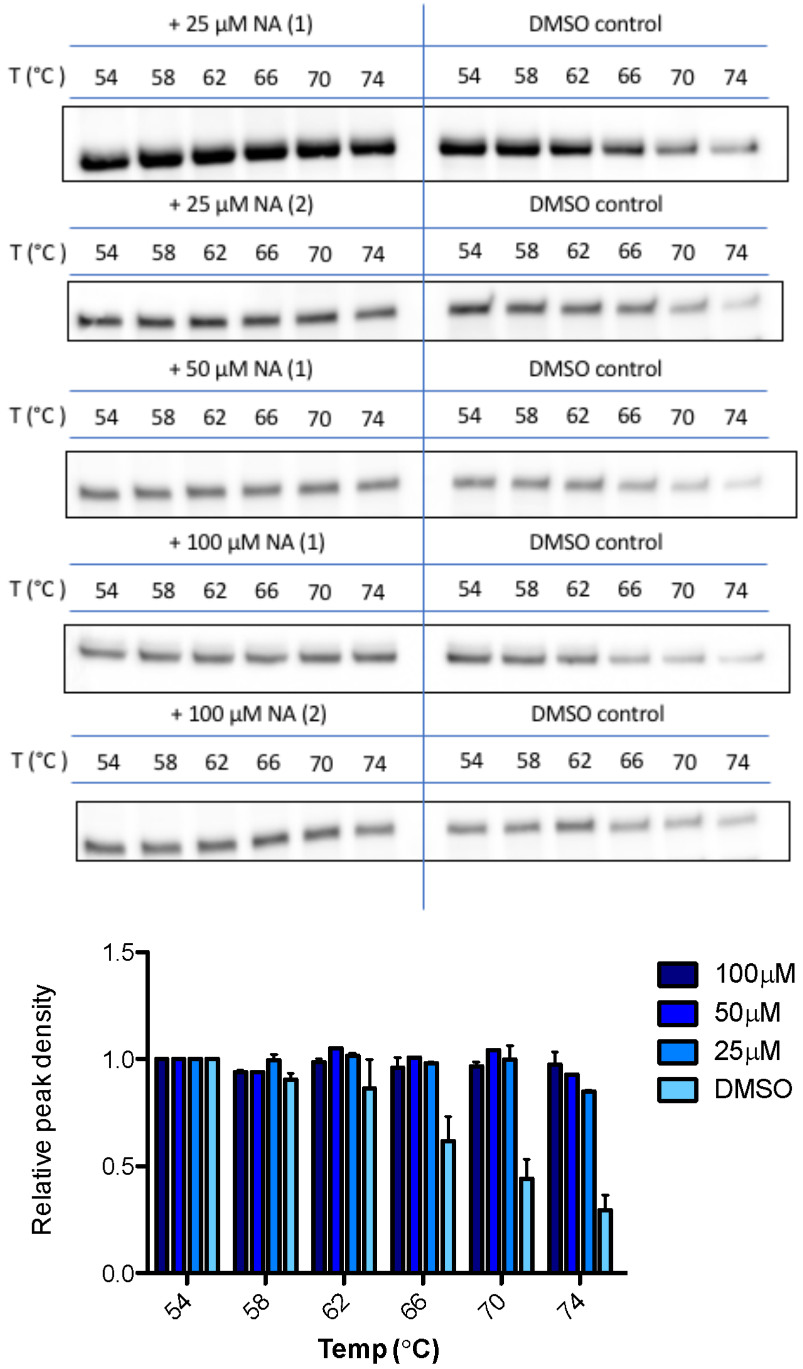


**Figure S10.** NA thermally stabilizes Med25 in cell lysates at multiple concentrations. Experiments were conducted at 25 µM (biological duplicate), 50 µM, and 100 µM (biological duplicate) NA. Equivalent DMSO was added to controls. Western blots for Med25, show thermal stabilization with NA for all conditions (top). Band intensities, calculated using ImageJ, at each temperature emphasize stabilization, showing only slight decrease in band intensity at 74C, 25 µM condition (bottom). Error bars represent the standard deviation of the mean from duplicate experiments.

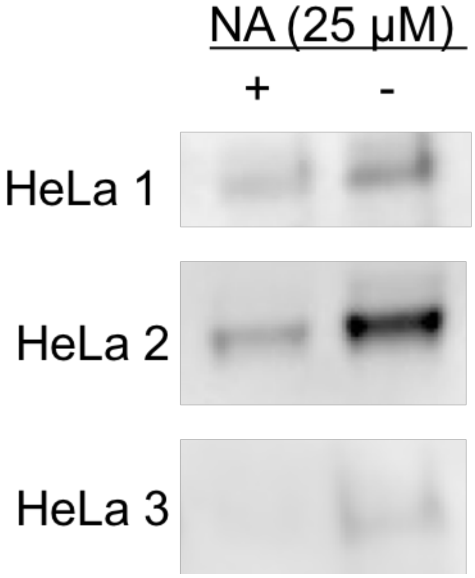


**Figure S11**. Replicate blots showing NA inhibition of co-immunoprecipitation of ETV5 with Med25 in HeLA cells. Blots show ETV5 protein identified via ETV5 antibody and visualized using chemilluminescense.

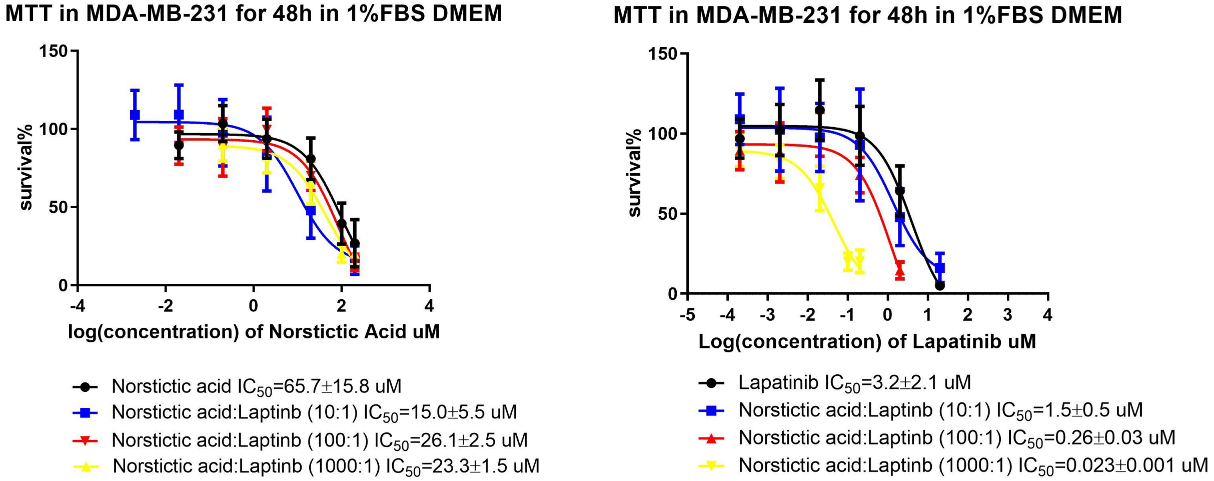


**Figure S12**. Dose-effect curves looking at MDA-MB-231 cell viability dosing with NA (left) and Laptinb (right) in various combinations. Error bars indicate the error compounded from one standard deviation of experiments performed in triplicate.

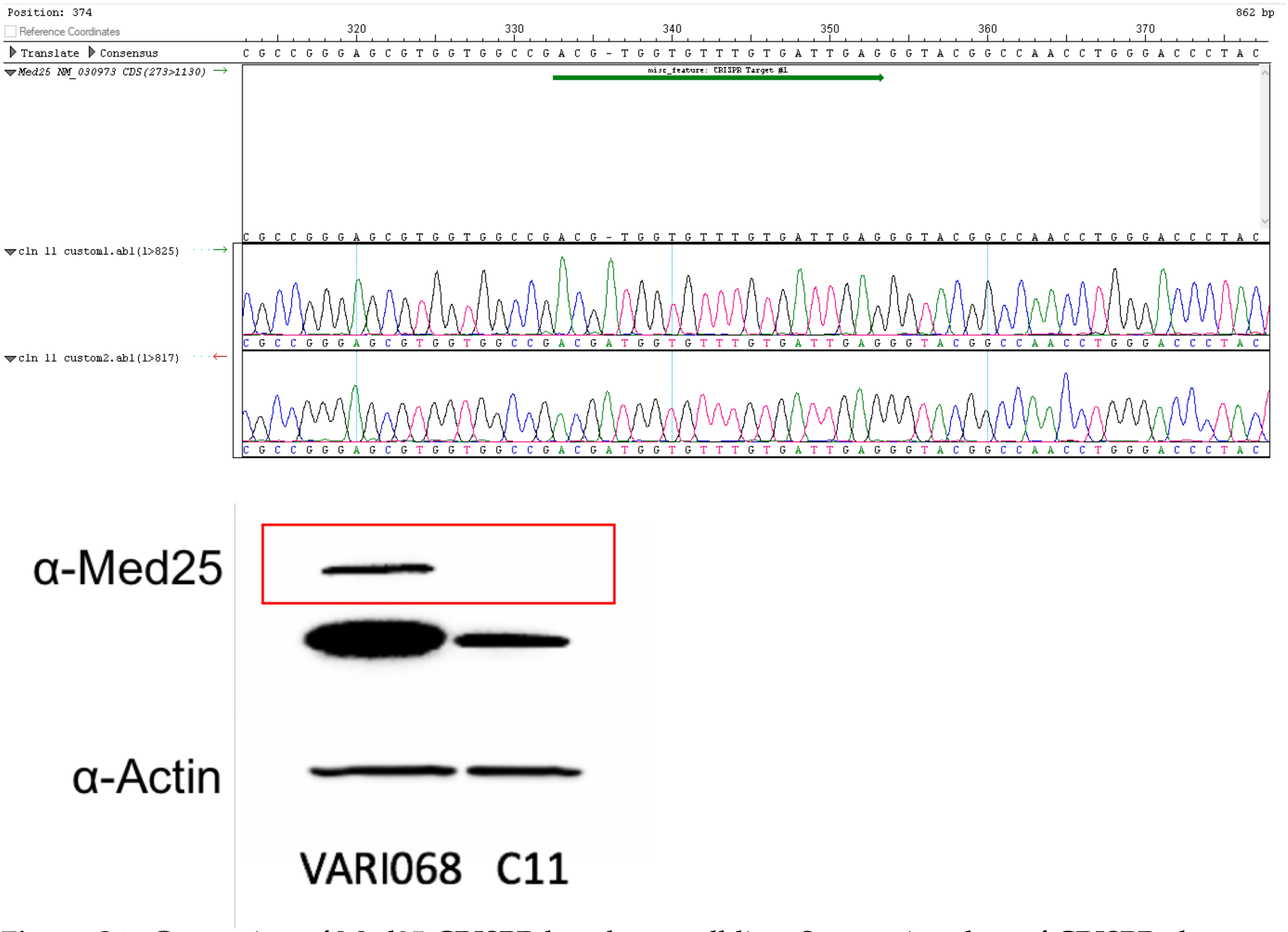


**Figure S13** Generation of Med25 CRISPR knock out cell line. Sequencing data of CRISPR clone 11 shows a 1bp insertion (adenosine) after guanidine 335, leading to knock out due to a frameshift mutation (top). Western blot with Med25 antibody shows absence of band corresponding to Med25 in CRISPR clone 11, validating the absence of Med25 protein.

**Protein expression and purification**
The Med25 expression plasmid, referred to as pAcID(394-543)-His6, was generously provided by Prof. Patrick Cramer.[1] pAcID(394-543)-His6 mutants, with the exception of K420R, were prepared using site directed mutagenesis with the primers listed below. Plasmid sequence identity was confirmed via standard Sanger sequencing methods at the University of Michigan DNA Sequencing Core and analyzed using SeqMan Pro from the Lasergene DNASTAR software suite.

Primers used for SDM:

pAcID-(K518R)-His6
F Pr. TCATGCTCCTGTACTCGTCCAGGAAGAAGATCTTCATGGGCCTCATCCC
R Pr. GGGATGAGGCCCATGAAGATCTTCTTCCTGGACGAGTACAGGAGCATGA
pAcID-(K519R)-His6
F Pr. TCATGCTCCTGTACTCGTCCAAGAGGAAGATCTTCATGGGCCTCATCCC
R Pr. GGGATGAGGCCCATGAAGATCTTCCTCTTGGACGAGTACAGGAGCATGA
pAcID-(K520R)-His6
F Pr. TCATGCTCCTGTACTCGTCCAAGAAGAGGATCTTCATGGGCCTCATCCC
R Pr. GGGATGAGGCCCATGAAGATCCTCTTCTTGGACGAGTACAGGAGCATGA
pAcID-(KK518RR)-His6
F Pr. TCATGCTCCTGTACTCGTCCAGGAGGAAGATCTTCATGGGCCTCATCCC
R Pr. GGGATGAGGCCCATGAAGATCTTCCTCCTGGACGAGTACAGGAGCATGA
pAcID-(KKK518RRR)-His6
F Pr. TCATGCTCCTGTACTCGTCCAGGAGGAGGATCTTCATGGGCCTCATCCC
R Pr. GGGATGAGGCCCATGAAGATCCTCCTCCTGGACGAGTACAGGAGCATGA
pAcID-(K411R/K413R)-His6
F Pr. GGGGTCCTGGAGTGGCAAGAGAGACCCAGACCTGCCTCAGTGGATGCCAAC
R Pr. GTTGGCATCCACTGAGGCAGGTCTGGGTCTCTCTTGCCACTCCAGGACCCC

Mutant K420R was made using PCR-driven overlap extension.[2] The flanking primers A and D were designed to extend into the pET21b vector. Primers B and C contain the mutated codon. The primer designs are as follows:

Primer A: 5′-GAA GGA GAT ATA CAT ATG TCA GTC TCC AAT AAG-3′
Primer B: 5′-GAT GCC AAC ACC CGA CTG ACG CGG TCA CTG-3′
Primer C: 5′ CAG TGA CCG CGT CAG TCG GGT GTT GGC ATC-3′
Primer D: 5′-GTG GTG GTG CTC GAG GTT GGT GAT GAC-3′

WT Med25 and mutants were expressed and purified from heat-shock competent Rosetta pLysS cells (Novagen), in Terrific Broth (TB) containing 0.1 mg/mL ampicillin and 0.034 mg/mL chloramphenicol, using previously described conditions.[1,3] Cells were grown at 37 °C to an optical density (OD600nm) of 0.8. Temperature was reduced to 18°C and protein expression was induced upon addition of IPTG to a final concentration of 0.5 mM. Post-induction, cells were incubated 16 hours at 18°C. Cells were pelleted via centrifugation at 6000xg for 20 mins at 4°C. Cell pellets were stored at -80°C prior to purification. The harvested pellet was thawed on ice and resuspended in 20 mL of lysis buffer (50 mM phosphate, 300 mM sodium chloride, 10 mM imidazole, pH 6.8). Cells were then lysed by sonication on ice and cellular lysates were cleared by centrifugation at 9500 rpm for 20 min at 4°C. The supernatant lysate was then added to 750 µL Ni-NTA beads (Qiagen) and incubated for 1 hour at 4°C. The resin was pelleted by centrifugation at 2500 rpm for 2 min at 4°C and washed with wash buffer (50 mM phosphate, 300 mM sodium chloride, 30 mM imidazole, pH 6.8) a total of five times. Protein was then eluted with 2 mL of elution buffer (50 mM phosphate, 300 mM sodium chloride, 400 mM imidazole, pH 6.8) a total of three times. Eluent was then pooled and purified by cation exchange FPLC (Source 15S, GE Healthcare) using a gradient of Buffer B (50 mM phosphate, 100 mM NaCl, 1 mM DTT, pH 6.8) in Buffer A (50 mM phosphate, 1 mM DTT). The FPLC purified protein was then dialyzed into storage buffer (10 mM phosphate, 50 mM NaCl, 10% v/v glycerol, 0.001% v/v NP-40, pH 6.8) overnight, concentrated, aliquoted, and stored at -80°C. Final protein was greater than 90% pure as determined by Coomassie stained polyacrylamide gel. Protein concentration was determined by UV-Vis spectroscopy using an extinction coefficient, $\varepsilon = 22{,}460\ M^{-1}\ cm^{-1}$.

CBP KIX (586-672) was expressed in BL21 DE3 *E. coli* as previously described.[4] Cells were grown to an optical density (OD 600nm) of 0.8 (37°C, 250 rpm), induced with 0.25 mM isopropyl b-d-1-thiogalactopyranoside (IPTG) for 16 hours at 20°C, harvested by centrifugation (20 min, 6500xg) and stored at -80°C. Cell pellets were lysed via sonication in lysis buffer (10 mM phosphate, 300 mM NaCl, 10 mM imidazole, pH 7.2) containing 2-mercaptoethanol and cOmplete protease inhibitor (Roche, 11873580001). The Hisx6 tagged protein was affinity purified using immobilized metal ion affinity chromatography (IMAC) on a HisTrap HP Ni Sepharose column (GE Healthcare). Elution was conducted using an imidazole gradient of 10 mM to 600 mM imidazole. An additional round of purification was completed using ion-exchange chromatography on a Source S column (GE Healthcare) in phosphate buffer (50 mM, pH 7.2) by eluting with a NaCl gradient from 0 to 1M. Purified protein was buffer-exchanged by dialysis (overnight, 4°C) into 10 mM phosphate, 100 mM NaCl, 10% glycerol, pH 6.8. Purity was determined by Coomassie stained polyacrylamide gel. Protein concentration was determined by UV-Vis spectroscopy using an extinction coefficient, $\varepsilon = 12{,}950\ M^{-1}\ cm^{-1}$. Purified protein samples (>90% pure) were aliquoted and stored at -80°C.

Med15 (1-345) was expressed and purified by as previously described.[5]

The expression plasmid for p300 TAZ1(324-423) was generously provided by Prof. Paramjit Arora.[6] Protein was expressed in BL21 DE3 *E. coli*. Cells were grown in LB containing 0.1 mg/mL ampicillin and 1 mM $ZnCl_2$ to an optical density (OD 600nm) of 0.8 (37°C, 250 rpm), cooled to 22°C and induced with 100 μM IPTC for 5 hours. Cells were harvested by centrifugation (20 min, 6500xg) and stored at -80°C. Cell pellets were lysed by sonication in lysis buffer (50 mM Tris, 150 mM NaCl, pH 6.3) containing cOmplete protease inhibitor (Roche, 11873580001). The GST tagged protein was affinity purified using a GSTrap column (GE Healthcare). After initial binding of the protein to the column, elution was conducted using a buffer containing 10 mM reduced glutathione. An additional round of purification was completed using ion-exchange chromatography on a Source S column (GE Healthcare) in phosphate buffer (50 mM, 1 mM DTT, pH 7.2) by eluting with a NaCl gradient from 0 to 1M. Purified protein was buffer-exchanged by dialysis (overnight, 4°C) into 10 mM Tris, 100 mM $NaH_2PO_4$, 10% glycerol, 1mM DTT, 100 μM $ZnCl_2$, pH 6.8. Purity was determined by Coomassie stained polyacrylamide gel. Protein concentration was determined by UV-Vis spectroscopy using an extinction coefficient, $\varepsilon = 49{,}110\ M^{-1}\ cm^{-1}$. Purified protein samples (>90% pure) were aliquoted and stored at -80°C.

**Synthesis of transcriptional activation domain peptides**

The peptides listed below were prepared following standard FMOC solid-phase synthesis methods on a Liberty Blue Microwave Synthesizer (CEM). FMOC deprotections were completed by suspending the resin in 20% piperidine (ChemImpex) in DMF supplemented with 0.2 M Oxyma Pure (CEM) and irradiating under variable power to maintain a temperature of 90°C for 60 seconds. Coupling reactions were completed by combining the amino acid (5 eq relative to resin; CEM, ChemImpex, and NovaBiochem), diisopropylcarbodiimide (7 eq, ChemImpex), and Oxyma Pure (5 eq) in DMF and irradiating under variable power to maintain a temperature of 90°C for 4 minutes. The resin was rinsed four times with an excess of DMF between all deprotection and coupling steps. N-terminal addition of fluoresceine isothiocyanate (FITC) residue was conducted by adding 1.2 eq in 5% diisopropylethylamine in dimethyl formamide for

18 hours at RT. Peptides were deprotected and cleaved from the resin for 4 hours in 90% trifluoroacetic acid (TFA), 5% thioanisole, 3% ethanedithiol (EDT) and 2% anisole. Crude peptides were filtered to remove resin, dried under nitrogen stream, and precipitated from cold ether. Peptide suspensions were transferred to a 15 mL falcon tube, centrifuged at 4000 g for 5 minutes at 4°C, and ether decanted. Crude peptides were resuspended in 20-40% acetonitrile, purified via HPLC on an Agilent 1260 HPLC using a semi-prep C18 column (Phenomenex). Pure fractions were collected and lyophilized to afford pure peptides. Final purity was determined via analytical HPLC and identify was confirmed using mass spectrometry. Analytical spectra were obtained using an analytical C18 column (Phenomenex) on an Agilent 1260 HPLC. Mass spectra were obtained using an Agilent 6230 LC/TOF and an Agilent 6545 LC/Q-TOF.

VP16 (465-490) - Fluorescein isothiocyanate and β-alanine were coupled to the N-terminus to produce FITC-βA-YGALDMADFEFEQMFTDALGIDEYGG. A gradient of 10-40% acetonitrile over 30 min was used for HPLC purification.

ETV5 (38-68) - Fluorescein isothiocyanate and β-alanine were coupled to the N-terminus to produce FITC-βA-ALDMADFEFEQMFTDALG. A gradient of 10-40% acetonitrile over 30 min was used for HPLC purification

ATF6α (40-66) - Fluorescein isothiocyanate and β-alanine were coupled to the N-terminus to produce FITC-βA-DTDELQLEAANETYENNFDNLDFDLDLM. A gradient of 10-40% acetonitrile over 30 min was used for HPLC purification

MLL (840-858) – Fluorescein isothiocyanate and β-alanine were coupled to the N-terminus to produce FITC-βA- DCGNILPSDIMDFVLKNTP. A gradient of 10-40% acetonitrile over 30 min was used for HPLC purification

myb (291-316) - Fluorescein isothiocyanate and β-alanine were coupled to the N-terminus to produce FITC-βA-KEKRIKELELLLMSTENELKGQQVLP. A gradient of 10-40% acetonitrile over 30 min was used for HPLC purification

IBiD (2063–2111) – The N-terminus was acetylated to produce Ac-SPSALQDLLRTLKSPSSPQQQQQVLNILKSNPQLMAAFIKQRTAKYVAN. A gradient of 10-50% acetonitrile over 40 min was used for HPLC purification.

ACTR (1041-1088) – Fluorescein isothiocyanate and β-alanine were coupled to the N-terminus to produce FITC-βA- PSNLEGQSDERALLDQLHTLLSNTDATGLEEIDRALGIPELVNQGQAL. A gradient of 10-50% acetonitrile over 40 min was used for HPLC purification.

HIF1α (786-826) - Fluorescein isothiocyanate and β-alanine were coupled to the N-terminus to produce FITC-βA-SMDESGLPQLTSYDCEVNAPIQGSRNLLQGEELLRALDQVN. A gradient of 10-30% acetonitrile over 60 min was used for HPLC purification.

**Direct binding and competition experiments**

Direct binding and competition experiments were performed using fluorescence polarization. Low volume, non-binding, black 384-well plates (Corning) were used and fluorescence polarization was measured using a PHERAStar multi-mode plate reader with polarized excitation at 485 nm and emission intensity measured through a parallel and perpendicularly polarized 535 nm filters. Data was analyzed using GraphPad Prism 5.0. For direct binding experiments, a binding isotherm that accounts for ligand depletion (assuming a 1:1 binding model of peptide to ACID) was fit to the observed polarization values as a function of protein concentration to obtain the apparent equilibrium dissociation, $K_d$:

$$y = c + (b - c) \times \frac{(K_d + a + x) - \sqrt{(K_d + a + x)^2 - 4ax}}{2a}$$

"a" and "x" are the total concentrations of fluorescent peptide and protein, respectively, "y" is the observed anisotropy at a given protein concentration, "b" is the maximum observed anisotropy value, and "c" is the minimum observed anisotropy value. Each data point is an average of three independent experiments with the indicated error representing the standard deviation of the three replicates. For competition experiments, curves were fit with a non-linear regression using the built-in equation "log(inhibitor) vs response – variable slope" from which the $IC_{50}$ value was calculated

**High-throughput screening**
Assays were performed in a final volume of 20 $\mu$L in a low volume, non-binding, black 384-well plate (Corning) and read by plate reader (Pherastar) with polarized excitation at 485 nm and emission intensity measured through parallel and perpendicularly polarized 535 nm filters. Optimization of fluorescence polarization assay for high throughput was conducted by testing stability of the VP16(465-490)•AcID interaction (Kd) over time, with combinations of DMSO and NP-40 (Figure S1). The assay shows little variance in affinity over time, up to 20 hours as well as tolerance to DMSO (5%) and NP-40 (0.001%). 4046 compounds were tested from the MS Spectrum 2000, Focused Collections, and BioFocus NCC libraries, which include known bioactive molecules, secondary metabolites, natural products, and FDA approved drugs. 200 nL of each compound in DMSO was first plated, followed by addition of 10 $\mu$L FITC-VP16(465-490). The compounds were then tested for fluorescence quenching before 10 $\mu$L of Med25 AcID protein was added. Plates were incubated for thirty minutes at room temperature and read by plate reader, as described above with gain settings determined based on a well from columns 23-24 (tracer only). Final concentration of AcID protein was 850 nM, final concentration of FITC-VP16(465-490) was 20 nM, and compounds were assayed at a concentration of 20 $\mu$M with a final DMSO concentration of 1% v/v. Data was published to and analyzed using MScreen (http://mscreen.lsi.umich.edu).

The primary screening campaign had an average Z′ score of 0.87, indicating an excellent assay, and a 1.6% hit rate. For the purposes of this screen, a hit was defined as any compound that resulted in inhibition greater than three standard deviations above the negative control, which corresponded to approximately ten percent inhibition. Following the primary screen, hits were filtered and compounds with known chemically reactive properties as well as those compounds that demonstrated native fluorescence greater than ten percent of the fluorescence produced by the tracer were removed.

**Mass spectrometry analysis of covalent adducts**
Protein (Med25 WT and mutants) was diluted to a concentration of 20 $\mu$M using storage buffer (10 mM phosphate, 50 mM NaCl, 10% v/v glycerol, 0.001% v/v NP-40, pH 6.8). Norstictic acid was added to the diluted protein to a final concentration of 20 $\mu$M (1 equivalent). Samples were incubated for 30 minutes at room temperature with gentle mixing on an orbital shaker. Samples were analyzed by mass spectrometry using an Agilent QToF LC/MS equipped with a Poroshell 300SB C8 reverse-phased column using a gradient of 5-100% acetonitrile with 0.1% formic acid in water with 0.1% formic acid over five minutes. Analysis of data was completed using the Agilent Qualitative Analysis Program with background subtraction and deconvolution settings for an intact protein of 10,000- 30,000 Da.

**Molecular dynamics simulations**

Modeling was performed using the NMR structure of Med25 AcID (PDB 2xnf). The norstictic acid was parameterized using CGENFF, which was then covalently linked to Med25 K519 through a PATCH that was created in CHARMM, with the molecules oriented out in space to allow for full, unbiased exploration around the protein before binding. The system was solvated using TIP3P water molecules as well as 100 mM NaCl using the MMTSB toolset so that the linked complex was in a cubic box with a minimum cutoff distance being 10 Å from the box edges. Simulations were unbiased molecular dynamics simulations using the CHARMM36 and CGENFF force fields for 100 ns of sampling at 298 K after allowing for 2ns of equilibration of the system. The simulation was run in the NVT ensemble using the Langevin dynamics algorithm with a friction coefficient of 5 $ps^{-1}$. The SHAKE algorithm was used to fix bond lengths during simulations. PME and vswitch were used for nonbonded interactions with a 12 Å cutoff. All molecular dynamics simulations were performed using GPUs through the CHARMM compatible OpenMM interface. Five independent trials of simulations were performed for each molecule.

**Cell Culture**

HeLa cells were purchased from ATCC (HTB-55). MDA-MB-231 cells were purchased from ATCC. VARI068 cells were derived from a patient-derived xenograft orthotopically implanted in NSG mice first and subsequent passaged through nude mice. HeLa cells were grown in DMEM (Gibco, 11965-092) supplemented with 10% FBS (R&D, S11150). MDA-MB-231 cells were grown in DMEM (Gibco, 11965-092) with 10% FBS (R&D, S11150). VARI068 cells were grown in DMEM (Gibco, 11965-092) supplemented with 10% FBS (R&D, S11150), and 1X Antibiotic-Antimicotic. Both cell lines were grown at 37 °C with 5% CO2.

**Thermal shift assays**

HeLa cells were grown, counted, and harvested using standard protocols (300,000 cells per temperature). Pelleted cells were resuspended in 1 mL cold PBS and transfered to a 1.5 mL Eppendorf tube on ice. Cells were again pelleted by centrifugation at 1500 G for 3 minutes at 4 °C. Nuclear extracts were generated using a NE-PER kit (Thermo Scientific, 78833) and manufacturer's protocol. After isolating the nuclear extract, a buffer exchange into PBS was conducted using a Zeba Spin Desalting Column 7K MWCO (Thermo Scientific, 89882). Prepared nuclear extracts were split into 2 epitubes. Norstictic acid (dissolved in DMSO) was added to one tube and an equivalent volume of DMSO was added to the other. Final concentration of DMSO was 0.1% v/v. Dosed nuclear extracts were incubated at room temperature for 30 minutes. After incubation, samples were alliquoted into thin-walled PCR tubes (15 $\mu$l per tube, the equivalent of 300,000 cells per tube).

A Labnet Multigene OPTIMAX PCR was used to heat each sample for 3 minutes. Six temperature points were tested, 54, 58, 62, 66, 70, and 74°C. Contents were transferred to epitubes and centrifuged at 17000 g for 1 minute at 4 °C to remove precipitated proteins. Contents of each epitube were carefully transferred to a clean epitube, leaving precipitated protein behind. LDS loading dye was added and samples were boiled for 5 minutes at 95°C. 10 $\mu$L of each sample was loaded onto a 4-20% mini-PROTEAN TGX gel (BioRad, 4561096) gel was run at 180V for 45 minutes. Protein was transferred from gels to PVDF membrane using a BioRaD TransferBox Turbo following the standard protocols. Membrane was blocked for 1 hour at room temperature using Super Block (Thermo Scientific, 37515). Med25 antibody (Novus biologicals, NBP2-55868) was added to membrane (1:1000 dilution in Super Block) and incubated overnight at 4 °C with gentle shaking. After removal of primary antibody and three washes with TBST, Secondary antibody (Santa Cruz SC-2004, 1:20,000 in Super Block) was added to membrane and incubated at RT for 1hr with shaking. After removal of secondary antibody with three washes with TBTS, HRP substrate (Thermo Scientific, 34095) was added and after 1 minute Western blot was visualized using Chemiluminescence on an Azure Biosystems c600 imager. Analysis was conducted using ImageJ software.

**Co-Immunoprecipitation**
Med25 antibody (Santa Cruz, sc-393759) was chemically crosslinked to Dynabeads Protein G (Invitrogen, 10004D) using Bis(sulfosuccinimidyl) suberate ($BS^3$). Briefly, 20 $\mu$L Dynabeads Protein G were washed with 250 $\mu$L PBST 3 times. Med25 Santa Cruz antibody (24 $\mu$L, 4.8ug) in 400 $\mu$L PBST was added to beads and incubated on with rotation at 4 °C for 1hr. Antibody coupled beads were washed twice with conjugation buffer (20mM sodium phosphate, 150 mM NaCl, pH = 7.5), resuspended in 250 $\mu$L of 5mM $BS^3$ (Thermo Scientific, 21580), and incubated at RT with rotation. After 30 minutes, 12.5 $\mu$L quenching buffer (1M Tris HCl, pH 7.5) was added and beads were incubated an additional 15 min with rotation. Crosslinked beads were washed three times with PBST and immediately used or stored at 4°C in PBST and used within 24 hours.

Mammalian cells were harvested as described previously. Nuclear extracts were generated from mammalian cells using a NE-PER kit (Thermo Scientific, 78833). Extracts were buffer exchanged into PBS using Zeba Spin Desalting Column 7K MWCO (Thermo Scientific, 89882). Extracts were dosed with either Norstictic acid (dissolved in DMSO) or an equivalent volume of DMSO (0.1% v/v). After washing crosslinked beads with PBS, nuclear extracts were added and incubated for 3 hours at 4°C with rotation. Flowthrough was collected and saved, and beads were washed 3X with PBST. LDS sample buffer (Invitrogen, NP0007; 2X final concentration) was added to beads and incubated at 95C for 10 minutes to elute immunoprecipitated proteins. Samples were run on 4-20% mini-PROTEAN TGX gels (BioRad, 4561096). Transfer and blotting was conducted using standard protocols (see cellular thermal shift assay protocol). ETV5 antibody (Proteintech, 13011-1-AP) diluted 1:2000 in SuperBlock for use.

**Quantitative polymerase chain reaction**
For endogenous gene expression analysis, HeLa cells were seeded into a 24-well plate ($1x10^5$ cells/well) and allowed to adhere overnight. Media was removed and replaced with OptiMem media containing vehicle or compound delivered in DMSO (0.5% v/v) at the indicated concentrations. After 6 h, the media was removed and total RNA was isolated using RNeasy Plus RNA isolation kits (Qiagen) according to manufacturer's instructions. Each RNA sample was used to synthesize cDNA using iScript cDNA synthesis kits (Bio-Rad). RT-qPCR reactions were carried out in triplicate in an Applied Biosystems StepPlusOne instrument using SYBR green master mix and primers for:

human RPL19 Forward. 5′:ATGTATCACAGCCTGTACCTG:3′
human RPL19 Reverse. 5′:TTCTTGGTCTCTCTTCCTCCTTG:3′
MMP-2 Forward. 5′:CATTCCAGGCATCTGCGATGAG:3′
MMP-2 Reverse. 5′:AGCGAGTGGATGCCGCCTTTAA:3′

RT-qPCR analysis was carried out using the comparative CT Method (ΔΔCT Method) to estimate MMP-2 mRNA levels relative to the reference RPL19 mRNA levels. Experiments were conducted in biological duplicate, technical triplicate.

**Synergy experiments with lapatinib**
Cells were seed at 3000 cells per well in 96 well plates. After 24 hours, cells were adhered on the plate. Medium was changed from 10% FBS to 1% FBS and at the same time, appropriate amount of compound was added (DMSO 1%). After 24 hours, old medium was removed and new 1% FBS DMEM medium and compounds were added to each well. The day after this treatment, cell viability was measured by Cell Proliferation Kit (MTT) from Roche following the manufacturer's instructions.

Calculation of synergy: All calculations were performed in Graphpad Prism and Microsoft Excel 365. Isobolograms were generated based on the dose fractions calculated by the $IC_{50}$s of either Norstictic acid, Laptinb alone or combination of them in different ratios. In this case, dose fraction is defined as the $IC_{50}$ of one component (Norstictic acid or Laptinb) in a combination divided by the $IC_{50}$ of that component in isolation required to exert the same effect. The dose fractions of Norstictic acid and Laptinb represent the x/y coordinates on the isobologram. Combination index (CI) is the sum of two dose fractions. For single agent, CI is 1 and for combination, synergy effect is present when CI<1.

Dose fraction of Compound A for combination AB = $IC_{50}$(A in AB) / $IC_{50}$(A in isolation)
Dose fraction of Compound B for combination AB = $IC_{50}$(B in AB) / $IC_{50}$(B in isolation)
CI (combination index) = dose fraction A + dose fraction B.

**Generation of Med25 CRISPR KO cell lines**
VARI-068 cell lines were transfected using the Nucleofector II system (Lonza) with pSpCas9(BB)-2A-GFP (PX458) (Posor et al., 2013), a gift from Dr Feng Zhang (Addgene, plasmid #48138), containing the target sequence 5′-CTCAATCACAAACACCACGT-′3 against Med25. Two days after transfection, single cells were sorted for GFP expression into 96-well plates. Following clonal expansion, genomic DNA was isolated and clones were screened for Med25 mutations using SURVEYOR reactions (IDT) with the primer pairs listed below. Positive clones were sequenced to identify specific mutational events and western blotted for Med25 (Novus biologicals, NBP2-55868). KO validation is shown in Figure S13.

Forward: 5′-GACTGAGCCGCTTTCAATTTAT-′ 3
Reverse: 5′-CTCAGCTCCTCCTTTCTCAGAC-′3

**Analytical HPLC traces of synthetic peptides**

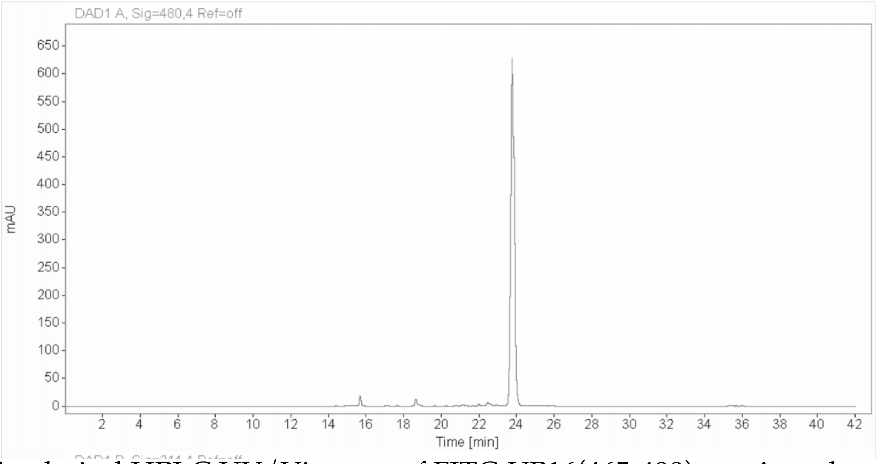


Analytical HPLC UV/Vis trace of FITC-VP16(465-490) monitored at 480 nm.

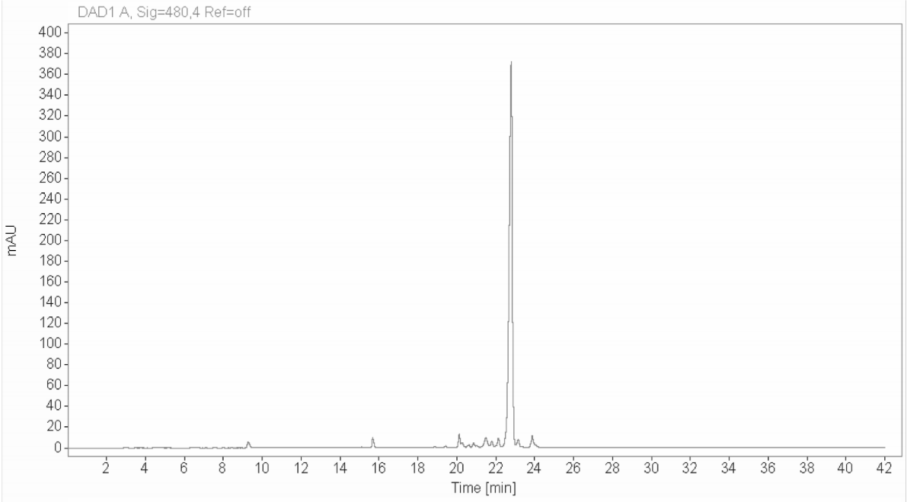


Analytical HPLC UV/Vis trace of FITC-ETV5 (38-68) monitored at 480 nm

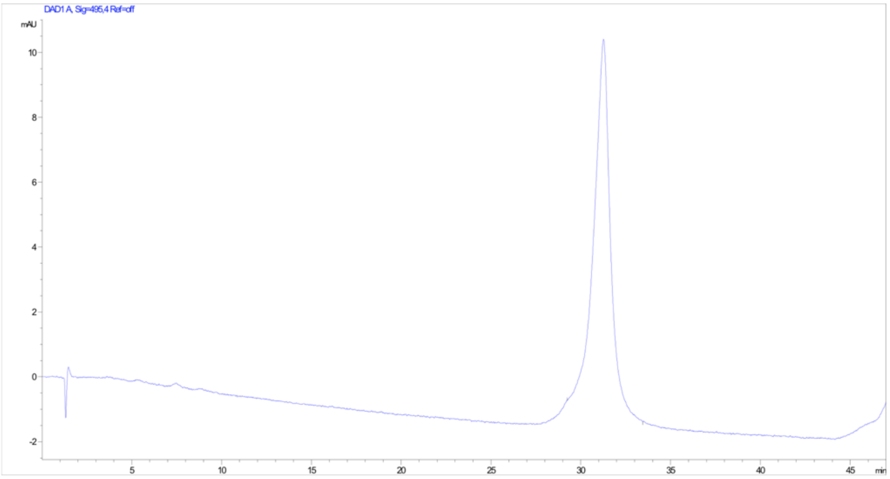


Analytical HPLC UV/Vis trace of FITC-ATF6α (40-66) monitored at 495 nm

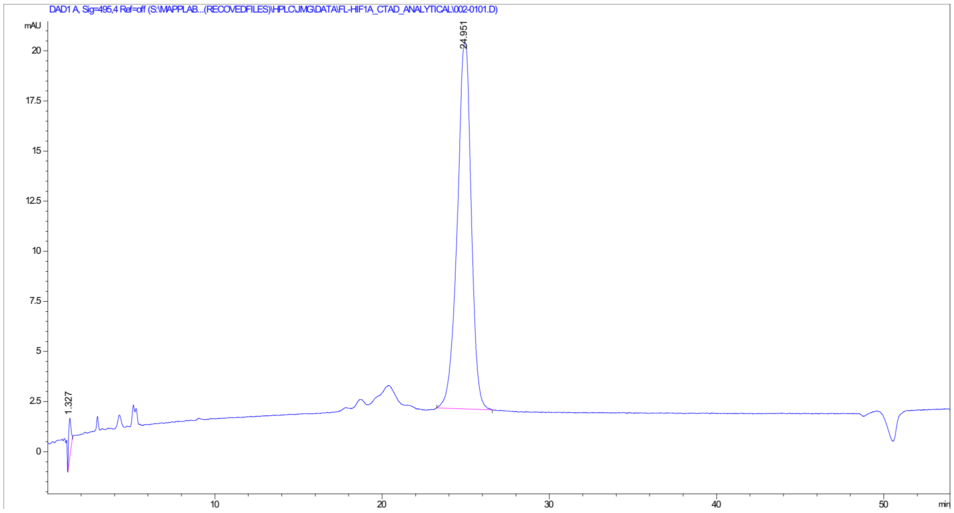

Analytical HPLC UV/Vis trace of FITC-HIF1α (786-826) monitored at 495 nm

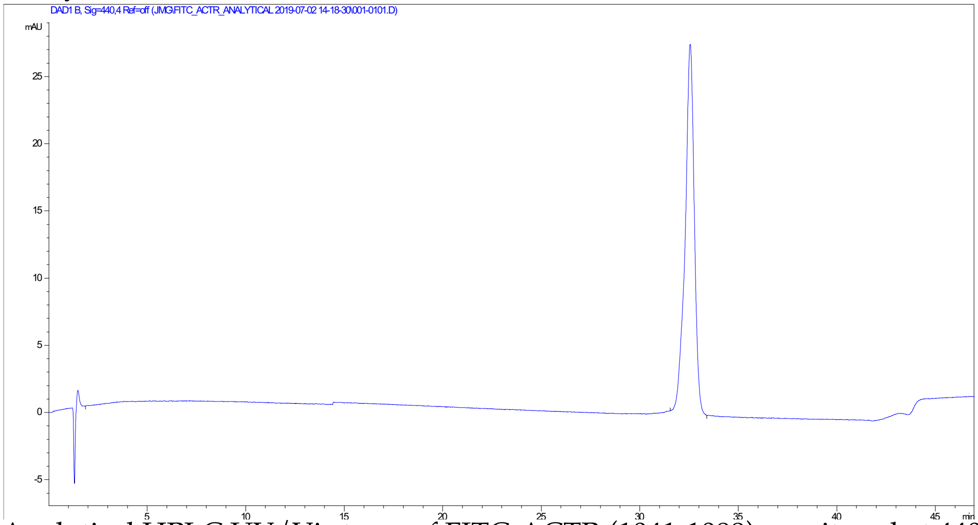

Analytical HPLC UV/Vis trace of FITC-ACTR (1041-1088) monitored at 440 nm.

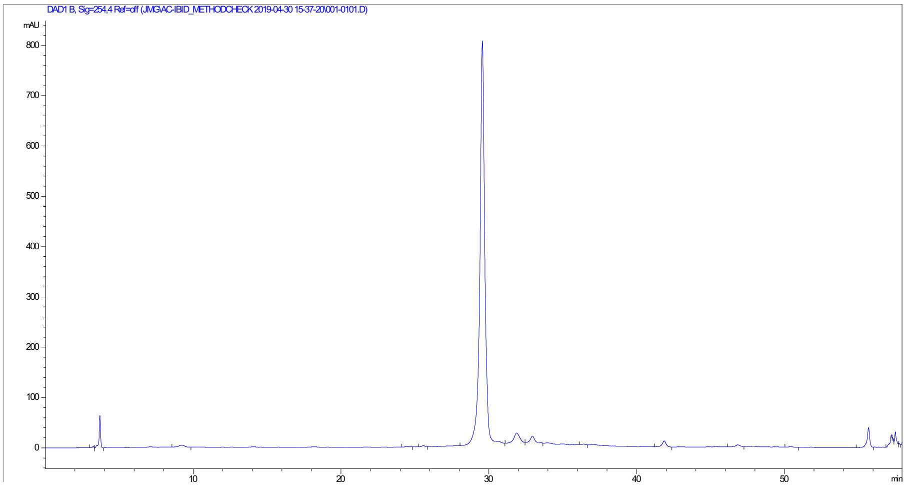

Analytical HPLC UV/Vis trace of Ac-IBiD (2063–2111) monitored at 254 nm.

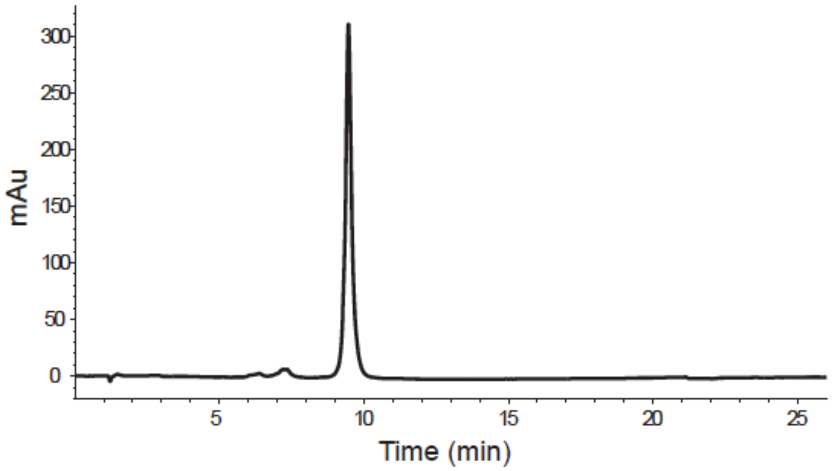

Analytical HPLC UV/Vis trace of FITC- Myb (291-316) monitored at 425 nm.

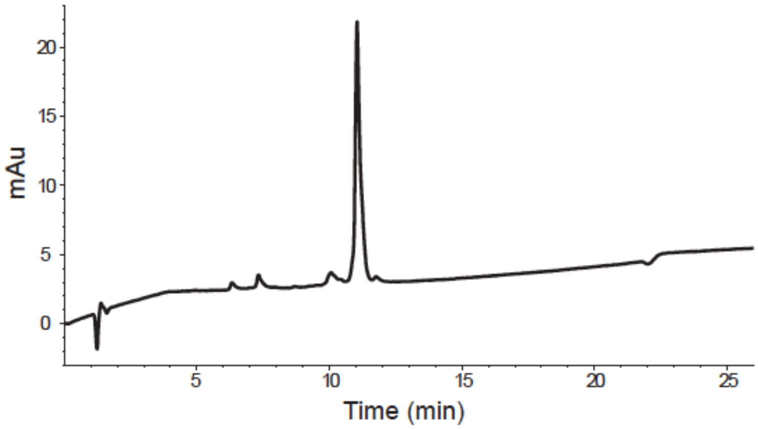


Analytical HPLC UV/Vis trace of FITC-MLL(840-858) monitored at 425 nm.
